# Circulating CD4^+^ memory T cells give rise to a CD69^+^ resident memory T cell population in non-inflamed human skin

**DOI:** 10.1101/490094

**Authors:** Maria M. Klicznik, Ariane Benedetti, Angelika Stoecklinger, Daniel J. Campbell, Iris K. Gratz

**Affiliations:** Department of Biosciences, University of Salzburg, Salzburg, Austria; Benaroya Research Institute, 1201 9th AVE, Seattle, WA 98101 USA; Department of Immunology, University of Washington School of Medicine, Seattle WA 98109, USA; Division of Molecular Dermatology and EB House Austria, Department of Dermatology, Paracelsus Medical University Salzburg, Austria

## Abstract

The blood of human adults contains a pool of circulating CD4^+^ memory T cells and normal human skin contains a CD4^+^CD69^+^ memory T cell population that produce IL17 in response to *Candida albicans*. Here we studied the generation of CD4^+^CD69^+^ memory T cells in human skin from a pool of circulating CD4^+^ memory T cells.

Using adoptive transfer of human PBMC into a skin-humanized mouse model we discovered the generation of CD4^+^CD69^+^ resident memory T cells in human skin in absence of infection or inflammation. These CD4^+^CD69^+^ resident memory T cells were activated and displayed heightened effector function in response to *Candida albicans*. These studies demonstrate that a CD4^+^CD69^+^ T cell population can be established in human skin from a pool of circulating CD4^+^ memory T cells in absence of infection/inflammation. The described process might be a novel way to spread antigen-specific immunity at large barrier sites even in absence of infection or inflammation.

---

Healthy human skin is protected by at least four phenotypically and functionally distinct subsets of memory T cells that are either recirculating or tissue-resident (Watanabe et al. 2015). Tissue resident memory T cells (T_RM_) are maintained within the tissue and show superior effector function over circulating memory T cells. T_RM_ are defined by the expression of CD69 and/or CD103, both of which contribute to tissue retention (Mackay et al. 2015; Mackay et al. 2013). CD4^+^ CD69^+^ T_RM_ are generated in response to microbes such as *C.albicans* and provide protective immunity upon secondary infection in mouse skin, and a similar CD4^+^CD69^+^ T cell population that produced IL17 in response to heat killed *C.albicans ex vivo* was found in human skin (Park et al. 2018). By contrast, a population of fast migrating CD69^-^ CD4^+^ memory T cells entered murine dermis even 45 days after *C.albcians* infection when the infection was already resolved, hence they were likely not recruited in response to antigen. These data suggest that in absence of infection/inflammation, T cell memory in human skin is heterogenous and composed of resident and migratory memory T cell populations. In mouse parabiosis experiments CD4^+^ memory T cells were able to modulate CD69 and/or CD103 expression, exit the skin and were found in equilibrium with the circulation. However, it is unclear if these cells could re-migrate to the skin and re-express CD69 upon entry into the tissue (Collins et al. 2016). Re-entry into the tissue to regain residency at distant skin sites would facilitate dispersion of protective immunity across large barrier organs such as the skin. The presence of a migratory memory T cell population in previously infected skin (Park et al. 2018) implies the existence of circulating memory T cells that can enter non-infected skin in steady state. So far it remains to be elucidated if human circulating CD4^+^ memory T cells can give rise to resident memory T cells in human skin in absence of acute infection.

We therefore set out to test whether human peripheral blood mononuclear cells (PBMC) contain a circulating population of memory CD4^+^ T cells with the potential to enter non-inflamed/non-infected skin sites distinct from the site of initial antigen-encounter, and assume the phenotype and function of CD69^+^CD4^+^ resident memory T cells upon tissue entry. For this, we utilized a novel xenografting mouse model (detailed in Klicznik et al. linked submission) in which we adoptively transferred human PBMC into immunodeficient NOD-*scid IL2rγ^null^* (NSG) mice (King et al. 2008) mice that carried engineered human skin (ES), which was devoid of any resident leukocytes. Within this model, designated huPBMC-ES-NSG, we found that transferred PBMC preferentially infiltrated the human but not murine skin, and that the human skin tissue promotes T cell maintenance and function in absence of inflammation or infection (Klicznik et al. linked submission). In line with previous reports (Clark et al. 2006), we found that human PBMC contained a proportion of skin homing CLA^+^CD45A^-^CD4^+^ memory T cells and, upon adoptive transfer skin-tropic CLA^+^CD4^+^ T cells were significantly enriched in the ES compared to the spleen (Fig. 1a). Interestingly among these CLA^+^ cells, the majority expressed the T_RM_ marker CD69 in the ES (Fig.1b). Importantly, the increased proportion of CD4^+^ T cells expressing CD69 in the ES was likely not due to tissue-derived inflammatory cues (Mackay et al. 2012) such as IL1α, IL1β, IL18, IL23, IFNγ, TNFα and TNFβ, which were found at equal or lower levels as in healthy human skin (Supplement Fig.1). Nor was CD69 expression due to local activation of T cells by antigen-presenting cells (APC), which are virtually absent in the ES of this PBMC-ES-NSG model (Klicznik et al., linked submission). Some of the CD69^+^ cells in the ES expressed CD103, a marker of human CD4^+^CLA^+^ T_RM_ (Klicznik et al. 2018) (Fig. 1c). Additionally, CD69^-^ but not CD69^+^ cells within the ES and the spleen expressed CD62L, a marker of circulating memory T cells (Watanabe et al. 2015) (Fig. 1d). These two distinct memory subsets, a resident memory-like CD4^+^ CD69^+^ and a CD69^-^CD62L^+^ migratory population, replicate two major sets of memory T cells in healthy human skin (Park et al. 2018). Taken together, these data show that circulating CD4^+^ memory T cells have the ability to up-regulate markers of tissue-residency upon entry into non-inflamed/non-infected human skin.

**Figure 1:**
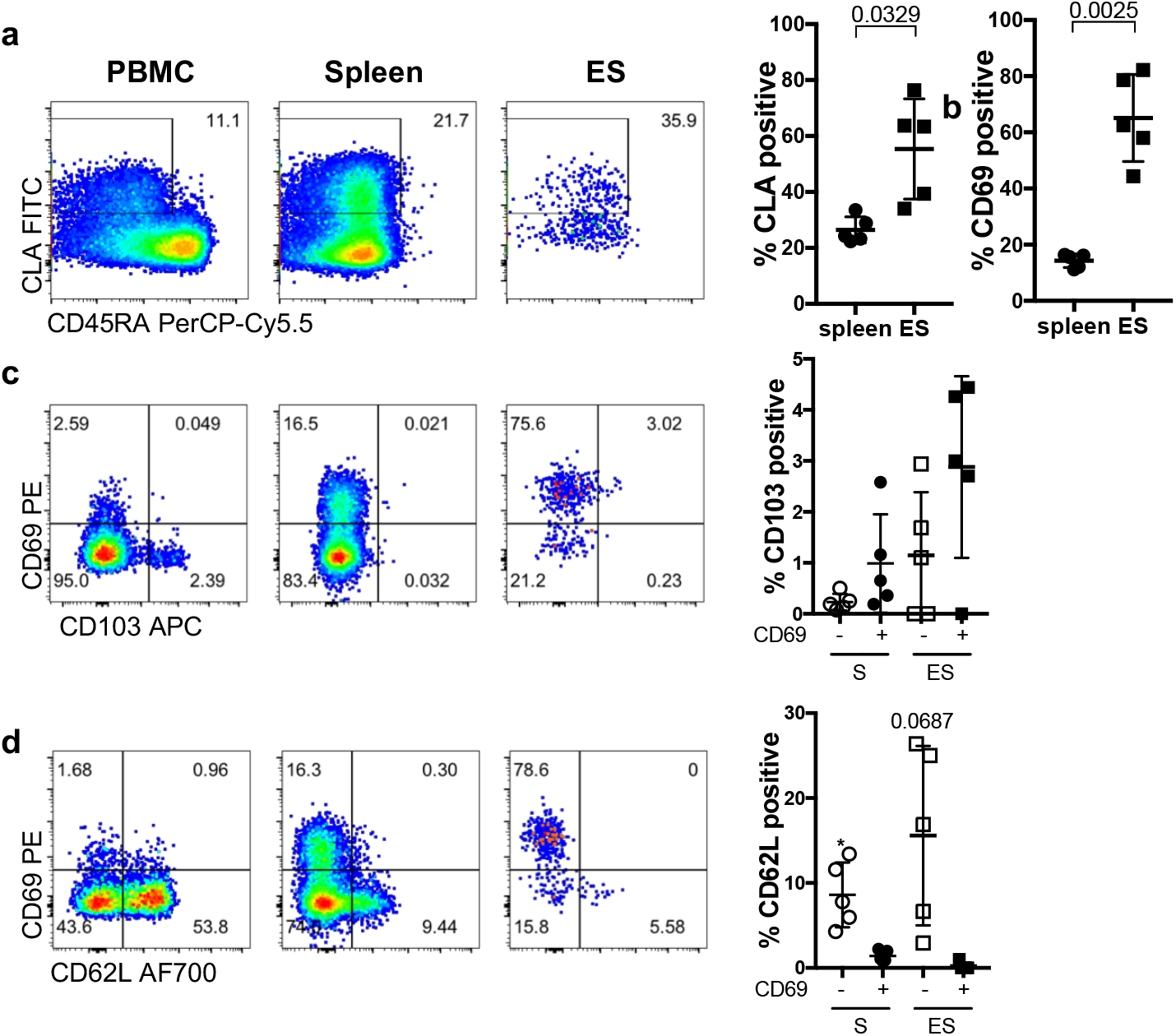
CD4^+^ memory T cells derived from the circulation up-regulate markers of residency and skin-tropism in engineered human skin. Engineered skin (ES) was generated on immunodeficient NSG mice. After complete wound healing (> 30 days) 3x106 autologous human PBMC were adoptively transferred. Single cell suspensions of spleen and ES of huPBMC-ES-NSG were prepared 21 days after adoptive transfer and analyzed by flow cytometry together with ingoing PBMC (**a-b**) Representative flow cytometry analysis and graphical summary of expression of indicated markers by CLA^+^CD45RA^-^CD4^+^CD3^+^ live leukocytes. Mean +/- SD, statistical significance determined by paired t-test; (**c-d**) Representative flow cytometry analysis and graphical summary of the expression of indicated markers by CD69^+^ or CD69^-^ cells in ES and spleen (S) gated on CLA^+^CD45RA^-^ CD4^+^CD3^+^ live leukocytes. Mean +/- SD; one-way RM ANOVA with Dunett’s test for multiple comparison to CD69^+^ cells in ES.

CD69^+^ T_RM_ represent a transcriptionally, phenotypically, and functionally distinct T cell subset at multiple barrier sites with enhanced capacity for the production of effector cytokines compared to circulating cells (Kumar et al. 2017). A proportion of skin-tropic human CD4^+^ T cells is specific for *C.albicans* (Acosta-Rodriguez et al. 2007), and to compare the function of CD69^+^ and CD69^-^ CD4^+^ T cells infiltrating the ES, we injected the ES after adoptive transfer of human PBMC with autologous monocyte derived DC that were pulsed with heat killed *C.albicans*. (Fig. 2a). Upon antigen-challenge CD69^+^ memory T cells produced higher levels of effector cytokines such as IL17, IFNγ, TNFα and IL2 (Fig. 2b-e), which is in line with a rapid memory response upon secondary infections.

**Figure 2:**
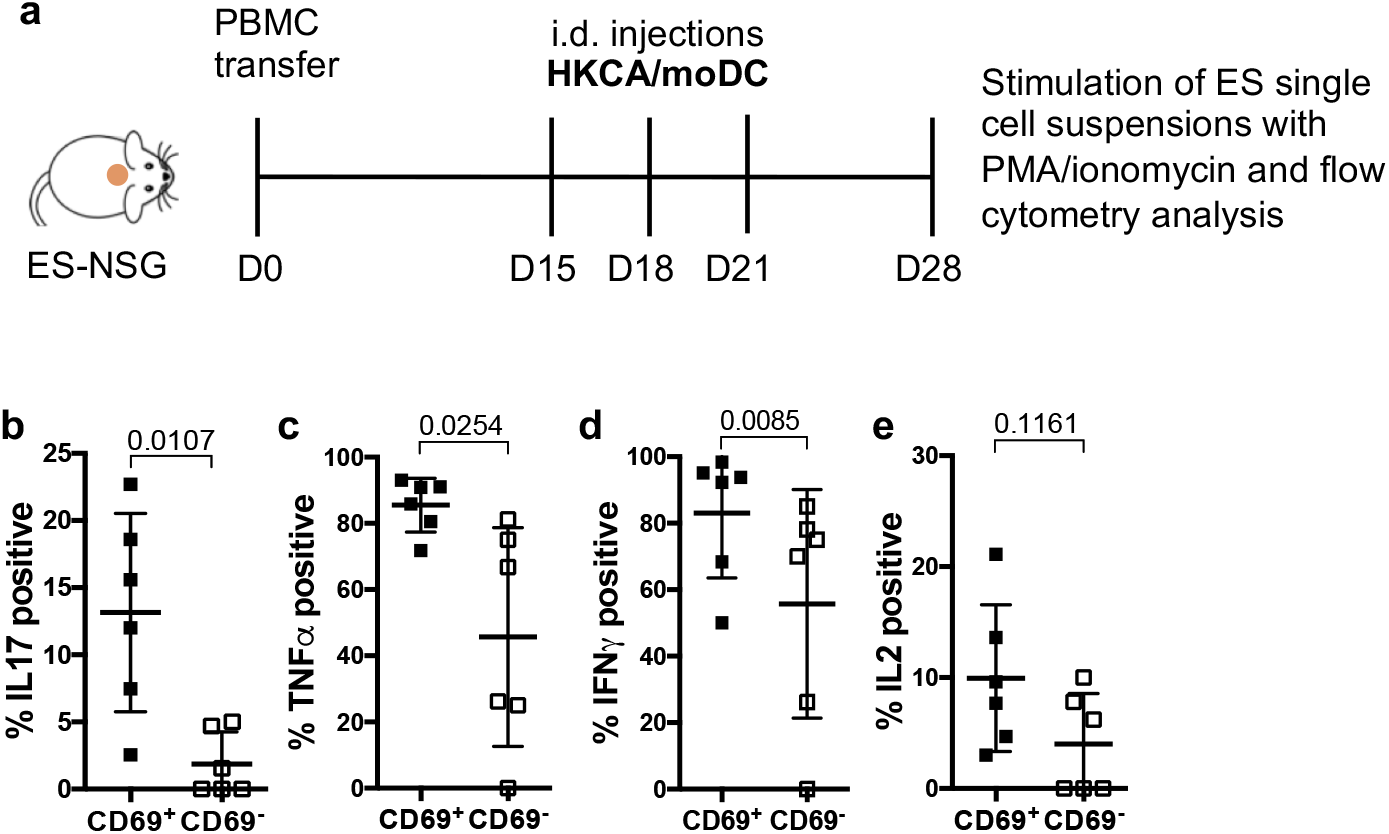
CD69^+^CD4^+^ cutaneous memory T cells show a distinct cytokine and activation profile in response to *C.albicans*. ES of huPBMC-ES-NSG mice was intradermally injected with heat killed *C.albicans* loaded onto autologous moDCs. Single cell suspensions of the ES were prepared and analyzed for intracellular cytokine secretion upon PMA/ionomycin stimulation (**a**) Experimental set-up; (**b-e**) Graphical summary of the proportion of positive cells of the indicated markers among CD69^+^ or CD69^-^ cells gated on CLA^+^CD45RA-CD4^+^CD3^+^ live leukocytes. Mean +/- SD; statistical significance determined by paired Student’s t-test

Here we show for the first time that circulating human memory CD4^+^ T cells can enter skin in the absence of infection or inflammation and give rise to a CD69^+^CD4^+^ memory T cell population (Fig.1b). Importantly these recently immigrated resident memory T cells produce substantially higher levels of effector cytokines (Fig.2b-e) in response to *C.albicans*, indicating that they represent the main responding memory population, which is in line with recent findings in murine skin (Park et al. 2018). It remains to be elucidated whether CD69^+^CD4^+^ memory T cells reside within the ES long-term or if they can modulate CD69 expression and leave the tissue again. The ability of circulating CD4^+^ memory T cells to enter non-inflamed skin sites, where they assume the phenotypical and functional profile of resident memory T cells reveals a novel mechanism to generate and disperse protective memory in large barrier organs, such as the skin.

## Acknowledgements

We thank all human subjects for blood and skin donation. We thank Dr. Stefan Hainzl, EB House Austria, Department of Dermatology, University Hospital of the Paracelsus Medical
University Salzburg, Austria, for the immortalization of primary human keratinocytes and fibroblasts. This work was supported by the Focus Program “ACBN” of the University of Salzburg, Austria, and NIH grant R01AI127726 awarded to IKG and DJC. MMK is part of the PhD program Immunity in Cancer and Allergy, funded by the Austrian Science Fund (FWF, grant W 1213) and was recipient of a DOC Fellowship of the Austrian Academy of Sciences.

## Author Contributions

IGK, DJC and MMK conceptualized the study, MMK designed and performed the experiments, MMK and AB acquired the data, GS acquired human samples, MMK performed data analysis, MMK and IGK interpreted data and wrote the manuscript. All authors reviewed the final version of the manuscript.

---

**Supplementary Figure 1:**
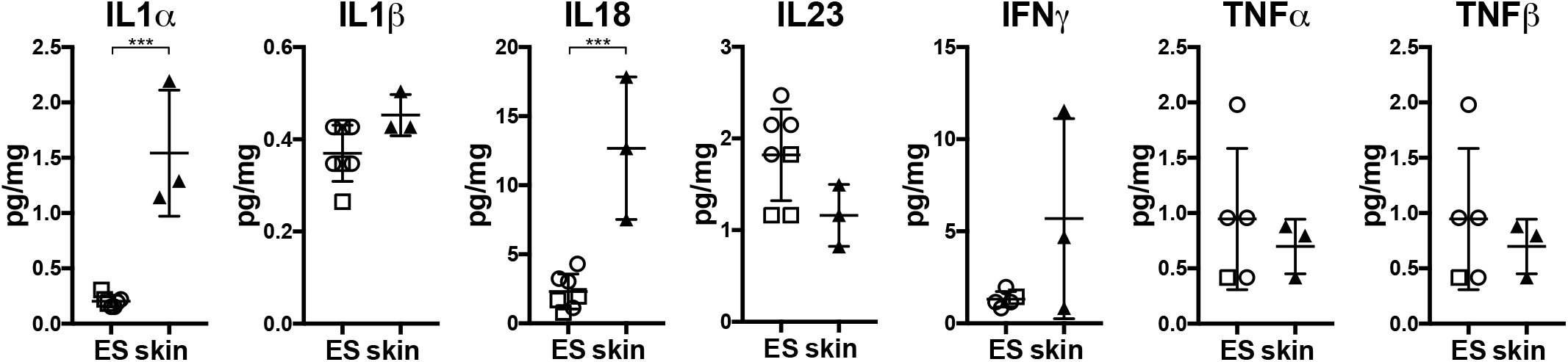
Analysis of pro-inflammatory cytokine levels in ES of huPBMC-ES-NSG mice in comparison to healthy human skin. ES and skin were weighed and analyzed for indicated cytokine levels using a bead-based multicomponent assay. Graphical summary shows concentration of indicated cytokine per mg skin; Mean +/- SD; statistical significance determined by paired Student’s t-test

## Material and Methods

### Mice

Animal studies were approved by the Austrian Federal Ministry of Science, Research and Economy. NOD.Cg-Prkdcscid Il2rgtm1Wjl/SzJ (NSG) mice were obtained from The Jackson Laboratory and bred and maintained in a specific pathogen-free facility in accordance with the guidelines of the Central Animal Facility of the University of Salzburg.

### Human specimens

Normal human skin was obtained from patients undergoing elective surgery, in which skin was discarded as a routine procedure. Blood and/or discarded healthy skin was donated upon written informed consent at the University Hospital Salzburg, Austria.

### PBMC isolation for adoptive transfer into NSG recipients and flow cytometry

Human PBMC were isolated from full blood using Ficoll-Hypaque (GE-Healthcare; GE17-1440-02) gradient separation. PBMC were frozen in FBS with 10% DMSO (Sigma-Aldrich; D2650), and before adoptive transfer thawed and rested overnight at 37°C and 5% CO_2_ in RPMIc (RPMI 1640 (Gibco; 31870074) with 5% human serum (Sigma-Aldrich; H5667 or H4522), 1% penicillin/streptomycin (Sigma-Aldrich; P0781), 1% L-Glutamine (Gibco; A2916801), 1% NEAA (Gibco; 11140035), 1% Sodium-Pyruvate (Sigma-Aldrich; S8636) and 0.1% β-Mercaptoethanol (Gibco; 31350-010). Cells were washed with PBS and 3x106 PBMC/mouse intravenously injected. Murine neutrophils were depleted with mLy6G (Gr-1) antibody (BioXcell; BE0075) as described before (Racki et al. 2010).

### Generation of engineered skin (ES)

Human keratinocytes and fibroblasts were isolated from human skin and immortalized using human papilloma viral oncogenes E6/E7 HPV as previously described (Merkley et al. 2009). Cells were cultured in Epilife (Gibco, MEPICF500) or DMEM (Gibco; 11960-044) containing 2% L-Glutamine, 1% Pen/Strep, 10% FBS, respectively. Per mouse, 1-2x106 keratinocytes were mixed 1:1 with autologous fibroblasts in 400μl MEM (Gibco; 11380037) containing 1% FBS, 1% L-Glutamine and 1% NEAA for *in vivo* generation of engineered skin as described (Wang et al. 2000).

### T cell isolation from skin tissues for flow cytometry

Healthy human skin and ES were digested as previously described (Sanchez Rodriguez et al. 2014). Approximately 1cm2 of skin was digested overnight in 5%CO_2_ at 37°C with 3ml of digestion mix containing 0.8mg/ml Collagenase Type 4 (Worthington; #LS004186) and 0.02mg/ml DNase (Sigma-Aldrich; DN25) in RPMIc. ES were digested in 1ml of digestion mix. Samples were filtered, washed and stained for flow cytometry or stimulated for intracellular cytokine staining.

### Flow cytometry

Cells were stained in PBS for surface markers. For detection of intracellular cytokine production, spleen and skin single cell suspensions and PBMC were stimulated with 50 ng/ml PMA (Sigma-Aldrich; P8139) and 1 μg/ml Ionomycin (Sigma-Aldrich; I06434) with 10 μg/ml Brefeldin A (Sigma-Aldrich; B6542) for 3.5 hrs. For permeabilization and fixation Foxp3 staining kit (Invitrogen; 00-5523-00) were used. Data were acquired on BD FACS Canto II (BD Biosciences) or Cytoflex LS (Beckman Coulter) flow cytometers and analyzed using FlowJo software (Tree Star, Inc.) A detailed list of the used antibodies can be found in the Supplements.

### Statistical analysis

Statistical significance was calculated with Prism 7.0 software (GraphPad) by RM ANOVA with Dunett’s multiple comparisons test, or by un-paired student’s t-test as indicated. Error bars indicate mean +/- standard deviation.

---

**Table S1:** Detailed list of antibodies and reagents

**Tissue preparation**
| Reagent | Company | Catalog number |
| --- | --- | --- |
| Collagenase Type 4 | Worthington | LS004186 |
| DNAse | Sigma-Aldrich | DN25 |
| RPMI 1640 | Gibco | 31870074 |
| human serum | Sigma-Aldrich | H5667/H4522 |
| Penicillin/streptavidin | Sigma-Aldrich | P0781 |
| L-Glutamine | Gibco | A2916801 |
| NEAA | Gibco | 11140035 |
| Sodium-Pyruvat | Sigma-Aldrich | S8636 |
| β-Mercaptoethanol | Gibco | 31350-010 |
| PBS | Gibco | 14190169 |
| Ficoll Paque Plus | GE-Healthcare | GE17-1440-02 |

**Cellular activation**
| Reagent | Company | Catalog number |
| --- | --- | --- |
| Brefeldin A | Sigma-Aldrich | B6542 |
| Foxp3 / Transcription Factor Staining Buffer Set | Invitrogen | 00-5523-00 |
| Ionomycin | Sigma-Aldrich | I06434 |
| PMA | Sigma-Aldrich | P8139 |
| Cytokine/Chemokine/Growth Factor 45-Plex Human ProcartaPlex™ | Invitrogen | EPX-450-12171-901 |

**Skin cell culture and transplantation**
| Reagent | Company | Catalog number |
| --- | --- | --- |
| Epilife | Gibco | MEPICF500 |
| DMEM | Gibco | 11960-044 |
| MEM | Gibco | 11380037 |
| TrypLE express | Gibco | 12604021 |

**Tissue preparation from mice**
| Reagent | Company | Catalog number |
| --- | --- | --- |
| BD Pharm Lyse | BD | 555899 |

**Antibodies**
| Reagent | Company | Catalog number |
| --- | --- | --- |
| CLA FITC (HECA-452) | Biolegend | 321306 |
| CD1a FITC (HI149) | BioLegend | 300104 |
| CD3 BV421 (UCHT1) | BioLegend | 300434 |
| CD3 bv605 (SK7) | BioLegend | 344835 |
| CD4 PE-594 (RPA-T4) | BioLegend | 300548 |
| CD4 PE-Cy5.5 (RPA-T4) | BioLegend | 300510 |
| CD8 PE (OKT8) | eBioscience | 12-0086-42 |
| CD14 bv421 (M5E2) | BioLegend | 301830 |
| CD45 BV785 (HI30) | BioLegend | 304048 |
| CD45RAPerCP-Cy5.5 (HI100) | BioLegend | 304122 |
| CD62L AF700 (DREG-56) | Biolegend | 304820 |
| CD69 PE (FN50) | BioLegend | 310906 |
| CD69 PerCP/eFluor710 (FN50) | eBioscience | 46-0699-42 |
| CD86 PE (B7-2) | eBioscience | 12-0869-41 |
| CD103 APC (Ber-ACT8) | BioLegend | 350216 |
| CCR7 bv605 (G043H7) | BioLegend | 353224 |
| IFNγ PE-Cy7 (B27) | BioLegend | 506518 |
| IL-2 PE (MQ1-17H12) | eBioscience | 12-7029-42 |
| IL17A APC (BL168) | BioLegend | 512334 |
| TNFα FITC (Mab11) | BioLegend | 502915 |
| Fixable Viability Dye eFluor™ 780 | eBioscience | 65-0865-14 |
| mLy6G (Gr-1) InVivoMab RB6-8C5 | BioXcell | BE0075 |

## Abbreviations

T_RM_: tissue resident memory T cells
PBMC: peripheral blood mononuclear cells
NSG: NOD-*scid IL2rγ^null^* mice
ES: engineered human skin
huPBMC-ES-NSG: NSG mice with grafted human PBMC and ES

